# Mesohabenular glutamatergic regulation of the consequences of uncontrollable stress

**DOI:** 10.64898/2026.07.21.739922

**Authors:** Dillon J. McGovern, Melissa Deming, Hadley Mills, Hannah Polatsek, Michael V. Baratta, David H. Root

## Abstract

Perceived or volitional control over a stressor is a critical determinant of resilient versus susceptible behavioral phenotypes. Activation of the lateral habenula (LHb) is required for the behavioral changes or “learned helplessness” effects that follow inescapable, but not escapable, stress. However, the relevant LHb inputs that regulate its mediation of inescapable stress outcomes are not fully elucidated. We find that ventral tegmental area (VTA) glutamatergic axons in LHb are activated by aversive stimuli, and this activation was not altered by prior inescapable stress experience. Similarly, optogenetic activation of VTA glutamatergic axons in LHb resulted in c-Fos expression in LHb neurons that did not depend on a prior inescapable stress experience. Photoinhibition of VTA glutamatergic axons in LHb during each inescapable stress trial prevented social and nonsocial consequences of inescapable stress. We interpret these results such that VTA glutamatergic inputs to LHb report aversive stimuli to LHb, possibly related to current salient aversive experience without regard to prior stress history. Nevertheless, VTA glutamatergic input to the LHb plays an important role in regulating LHb-mediated social and nonsocial consequences of uncontrollable stress.

## 1.0 INTRODUCTION

The perceived degree of behavioral control over a stressor can be a potent modulator of its impact. The neural basis of stressor controllability has been studied in animals by comparing the effects of escapable stress (ES) with those of physically identical yoked inescapable stress (IS). Inescapable stress produces a constellation of behavioral changes that do not follow escapable stress, such as reduced sociability and exaggerated fear (Christianson et al., 2008; Maier and Seligman, 2016; Baratta et al., 2023). In mice, we have found that the behavioral consequences of inescapable stress are sex-specific. Males result in social avoidance and potentiated fear while females instead result in exploratory deficits within the light-dark box (McGovern et al., 2024), a task associated with anxiety-like behavior (Rosso et al., 2022). While many brain regions participate in anxiety-like or depression-like behaviors, lesions or chemogenetic silencing of the lateral habenula (LHb) block the development of IS-induced social avoidance and shuttlebox escape deficits (Amat et al., 2001; Dolzani et al., 2016), establishing its role as an important regulator of IS outcomes.

LHb is an evolutionarily conserved epithalamic structure critical for the integration of motivationally-relevant stimuli into actions via descending regulation of multiple monoaminergic systems (Lecca et al., 2014; Zahm and Root, 2017; Yang et al., 2018a). In nonhuman primates and humans, LHb activity is associated with negative affect and depression, or anxiety-like states (Matsumoto and Hikosaka, 2009; Savitz et al., 2011; Savitz et al., 2013; Hennigan et al., 2015) and deep brain stimulation targeting the epithalamus was efficacious in reducing depression scores in a case study (Sartorius et al., 2010). Consistent with these observations, rodent models demonstrate that LHb hyperactivity promotes depression-like and anxiety-like behavior (Yang et al., 2008; Li et al., 2011; Li et al., 2013; Shabel et al., 2014; Meye et al., 2015; Berger et al., 2018; Kang et al., 2018; Purvis et al., 2018; Yang et al., 2018b; Cerniauskas et al., 2019; Gold and Kadriu, 2019; Lecca et al., 2023). However, the afferent pathways that influence LHb’s mediation of these processes are not wholly understood.

The LHb receives diverse inputs from brain regions that play critical roles in the consequences of inescapable stress, such as the bed nucleus of the stria terminalis, amygdala, and ventral tegmental area (VTA) (Hammack et al., 2004; Christianson et al., 2010; Christianson et al., 2011; Meye et al., 2013; Zahm and Root, 2017). Within the VTA, neurons expressing vesicular glutamate transporter 2 (VGluT2) are required for the social and nonsocial consequences of inescapable stress and about one-third of LHb neurons receiving VTA glutamatergic inputs are c-Fos activated following inescapable stress (McGovern et al., 2024). Here, we examined whether the VTA glutamatergic pathway to LHb causally participates in the social and nonsocial consequences of inescapable stress. We found that VTA glutamatergic axons in LHb were activated by aversive stimuli, and that optogenetic activation of this LHb afferent results in c-Fos activity, but neither were affected by prior inescapable stress experience. Optogenetic inhibition of VTA glutamatergic or dual glutamatergic/GABAergic axons in LHb reduced social and nonsocial consequences of inescapable stress. We interpret these data such that the VTA→LHb glutamatergic pathway signals events related to current aversive experiences without regard to prior stress and is one necessary component of the LHb influence on the behavioral consequences of uncontrollable stress.

## 2.0 METHODS

### 2.01 Stereotactic surgery

Male and female VGluT2::Cre mice (n=79, 8-12 weeks of age) received viral injections into the VTA (AP: -3.20, ML: 0.0, DV: -4.33; 400 nL at 100 nL/min) using an UltraMicroPump (Micro4; World Precision Instruments) equipped with Nanofil syringes (World Precision Instruments, Sarasota, FL) and 35-gauge blunt needles. Following injection, the syringe remained in place for 10 minutes to minimize diffusion before a slow withdrawal. For *in vivo* monitoring of presynaptic calcium transients, mice were injected with AAV9-hSynapsin1-FLEX-axon-GCaMP6s (Addgene 112010; titer=9.67×10^12^). To measure GCaMP fluorescence, a unilateral optic fiber (400 μm core, 0.66 NA; Doric Lenses) was implanted dorsal to LHb (AP: - 1.45, ML: +0.30, DV: -2.8). For excitation experiments, mice were injected with AAV8-nEF-Con/Foff-ChRmine-oScarlet (Addgene 137161, titer=7×10^12^). For inhibition experiments, VGluT2::Cre mice were injected with AAV5-EF1a-DIO-eNpHR3.0-EYFP (Addgene 26966; titer=5×10^12^) or the control vector AAV8-hSyn-DIO-GFP (Addgene 50457; titer=11.5×10^12^). To inhibit glutamate and GABA co-transmitting inputs to LHb, VGluT2::Cre/VGaT::Flp mice (n=13) were injected with AAV8-nEF-Con/Fon-NpHR3.3-EYFP (Addgene 137152, titer=8×10^12^). To photoactivate halorhodopsin or ChRmine, bilateral optic fibers (200 μm core, 0.37 NA; Doric Lenses, Québec, Canada) were implanted dorsal to LHb (AP: -1.45 ML: ±1 at a 10-degree angle, DV: -2.7). All implants were secured with skull screws and dental cement (DuraLay, Alsip, IL). Mice were provided with 3 days of postoperative care, consisting of daily carprofen injections (5 mg/kg) followed by 3-4 weeks of recovery before experimentation.

### 2.02 Stressor Controllability Paradigm

Mice were randomly assigned to IS or no stress (NS) groups at surgery. A group of male wildtype C57Bl6/J mice underwent ES to provide the temporal pattern (shock onset/offset) and duration of tailshocks to IS mice. For the stress procedure, mice were placed in Plexiglas chambers (7 × 6 × 9 cm; Med Associates, St. Albans, VT) featuring a front-mounted wheel and a rear tail-restraint rod. The mouse’s tail was secured to the rod using surgical tape and prepared with electrode cream and two copper electrodes. Tail shocks were delivered to the copper electrodes via alligator clips connected to a shock generator (MED-Associates). While the chambers were physically identical, the wheel in the IS chamber was locked, whereas the ES wheel remained mobile to allow for shock termination. The ES wheel was further modified to produce an auditory/tactile click every quarter-turn to provide immediate response-related feedback. Shock delivery and behavioral criteria were managed using custom-written MATLAB code connected to a LabJack. The single shock session consisted of 100 trials of unsignaled tail shocks separated by a random intertrial interval (ITI) averaging 60 seconds. Shock intensity was calibrated based on animal bodyweight (≤30g = 0.3mA, 30-35g = 0.35mA, and ≥35g = 0.40mA). For ES mice, shock termination was contingent upon an incremental wheel-turn requirement that increased across successful trials according to the following sequence: 1, 2, 4, 8, and 12 quarter-turns of the wheel. The response requirement increased following successful trials and decreased following unsuccessful trials. Success was defined as meeting the turn criterion within 7.5 seconds of shock onset, while failure was defined as exceeding 25 seconds. If no response occurred within 30 seconds, the shock was terminated automatically, and the criterion was reset to 1 quarter-turn. To ensure identical stressor exposure between groups, male halorhodopsin IS mice were yoked to ES mice, receiving tail shocks of exact intensity, duration, and timing as an escapable stress. For experiments comparing IS and NS groups, IS was achieved by replaying the shock timings from previously performed ES animals while NS mice remained in the home cage. For the Axon GCaMP6s cohorts, NS mice served as environmental controls and were individually housed in a novel cage for the duration of the stress session. Following the stress session, all mice were individually housed. Post-stress single housing was implemented because prior work has demonstrated that group housing can attenuate or abolish behavioral consequences of IS, particularly in male mice, likely due to social buffering effects (Hennessy et al., 2009).

### 2.03 Optogenetic stimulation

For inhibition experiments, halorhodopsin was activated using 589 nm light (8-10 mW, continuous) between tailshock onset and tailshock offset during IS. For ChRmine stimulation experiments, mice were isolated within the outside chambers of a conditioned place preference insert (18cmx20cm, Condition Place Preference within AnyBox, Stoelting). One day prior to IS, mice were isolated into the outside chambers and allowed to habituate to the apparatus for 1 hour with no stimulation. The next day, mice received IS or NS as described. One day following IS or NS, ChRmine was activated using 589 nm light (8-10 mW) at 20 Hz with 10 ms pulse duration and for periods involving 5 seconds on and 25 seconds off over 15 minutes. Ninety minutes later mice were euthanized as described in *Histology 2.08*.

### 2.04 Calcium fiber photometry recordings and analysis

In axon-GCaMP experiments, we examined whether footshock-related activity was modulated by prior stress experience. Specifically, we recorded axonal activity while mice received five footshocks before and after IS or NS. Mice were brought to MED-Associates mouse chambers equipped with grid floors. After one minute, a total of five footshocks (0.5 mA, 500 ms) were delivered, each separated by one minute. One day later, mice received either inescapable stress or no stress as described. One day following IS or NS, mice were brought back to MED-Associates chambers and delivered five footshocks as described. During footshocks, Axon GCaMP6s was excited at two wavelengths, 465-and 405-nm isosbestic control, by amplitude modulated signals from two light-emitting diodes reflected off dichroic mirrors and then coupled into an optic fiber as previously described (McGovern et al., 2021). The Axon GCaMP6s signal and the isosbestic control signal were returned through the same optic fiber and acquired using a LUX Photosensor, digitized at 1kHz, and then recorded by a real-time signal processor (Tucker Davis Technologies). The isosbestic signal (405nm) and the Axon GCaMP6s signal (465nm) were down sampled (10x) and peri-event time histograms were created surrounding shock onset. For each trial, data were detrended by regressing the 405nm signal on the 465nm signal. The generated linear model was used to create a predicted 405nm signal that was subtracted from the 465nm signal to remove movement, photo-bleaching, and fiber bending artifacts. Trials were normalized by z-score where average and standard deviation of the z-scores taken from -3.1 to -0.1 prior to event onsets. Trials were averaged and baseline maximum z-scores were taken from -6 to -3 seconds prior to event onset. Shock maximum z-scores were taken from 0 to 2 seconds following footshock onset.

### 2.05 Three-chamber sociability test

Twenty-four hours after tailshock or no stress, social interaction levels were assessed using the three-chamber sociability test in male mice. The test consisted of placing an ES, IS, or no stress subject in the center connecting compartment of an arena with three interconnected compartments (ANY-box, Stoelting, Wood Dale, IL). In one side, an empty sociability cage (Stoelting) was placed in the center and the other side contained an identical sociability chamber with a novel same-sex conspecific (approximately two weeks younger than the test mouse). Video recordings were made of mouse exploration of the entire arena for 10 min (30 Hz, ANY-maze, Stoelting). The social exploration test was quantified by counting the time the mouse spent exploring the cages containing the social and non-social stimuli. Exploration was defined as direct contact with the cage, including sniffing and rearing onto the apparatus. Social scores were quantified by the following formula: (social interaction time – nonsocial interaction time) / (social interaction time + nonsocial interaction time). Scores closer to 1 indicate a more social behavioral phenotype and scores closer to 0 reflect social avoidance. Videos were scored by experimenters that were blind to experimental condition.

### 2.06 Shock-elicited freezing

Thirty minutes after the sociability paradigm, male mice were placed inside a novel chamber with grid flooring (Med Associates). The shape, size, flooring, and illumination of the chamber differed from the context in which prior tailshock occurred. Subjects were allowed to explore the chamber for 2 min before receiving two footshocks (0.5 s, 0.5 mA, 1-min interstimulus interval) delivered through the grid floor. Within each chamber a video camera (AnyMaze, Stoelting) was connected to a computer for offline analysis of freezing behavior time-locked to shock onset. Freezing was defined as the absence of movement except that required for respiration. Videos were scored by experimenters blind to condition.

### 2.07 Light/Dark Box

Twenty-four hours after tailshock, IS or NS female mice were placed in the center of the dark compartment of the light/dark box (Stoelting). The light compartment was illuminated at 600 lumens. Video recordings were made of mouse exploration of the entire arena for 5 min (30 Hz, ANY-maze). Exploratory behavior was quantified by the latency to exit the dark compartment as well as total time spent in the light compartment (ANY-maze). If an animal did not exit the dark compartment, then a maximum latency of 300 sec was scored.

### 2.08 Histology

Mice were deeply anesthetized with isoflurane and perfused transcardially with 0.1M phosphate buffer (PB) followed by 4% (w/v) paraformaldehyde in PB. Brains were extracted, post-fixed overnight in the same fixative and cryoprotected in 18% sucrose in PB at 4°C. Coronal sections containing the LHb and VTA (30 µm) were taken on a cryostat, mounted to gelatin-coated slides, and imaged for axon GCaMP6s, halorhodopsin, or ChRmine expression and optical fiber cannula placement on Zeiss Axioscope. Data from mice with optic fibers not localized to the LHb or lacking viral expression were not analyzed.

### 2.09 c-Fos immunolabeling

LHb sections were incubated with blocking solution (4% BSA in 0.1 M PB, pH 7.3, supplemented with 0.3% Triton X-100) for 1 h. Sections were then incubated with guinea pig anti-c-Fos (1:500 in blocking solution, Synaptic Systems 226308) overnight at 4°C. Sections were washed in PB and incubated for 2 h at room temperature with donkey anti-guinea pig Alexa647 (1:200 in blocking solution, Jackson ImmunoResearch), mounted on gelatin-coated slides, coverslipped, and imaged on a Nikon A1R confocal (20X). c-Fos-labeled LHb neurons (Alexa647) were counted in FIJI (Schindelin et al., 2012).

### 2.10 Statistical analysis

Axon GCaMP was analyzed with repeated measures ANOVA across 2 days (pre-stress, post-stress) and 2 epochs (baseline, footshock). Wheel-turn performance was analyzed with repeated measures ANOVA across ten blocks of ten trials. Dunnet adjusted posthoc tests compared the first block with the remaining blocks. Social investigation was analyzed with a mixed ANOVA within-subjects object (social, nonsocial) and between-subjects experimental group. Sidak-adjusted pairwise comparisons followed up a significant object x group interaction. Social score was calculated as: (Time with Social Object – Time with Nonsocial Object) / (Time with Social Object + Time with Nonsocial Object). Social score and time freezing were analyzed with one-way ANOVA. Main effects of group were followed up simple contrast tests versus the inescapable stress GFP control group. Mann-Whitney U tests were used to analyze c-Fos between IS and NS groups as well as latency to enter the light side and time in light from light-dark box tests.

## 3.0 Results

### 3.1 VTA glutamatergic axons in LHb are activated by aversive events

We first assessed the sensitivity of VTA glutamatergic axons in LHb to aversive stimuli in mice with or without prior inescapable stress experience (**Figure 1A**). VGluT2::Cre mice were injected in VTA with AAVs encoding Cre-dependent Axon-localized GCaMP6s and an optic fiber was implanted dorsal to LHb. VTA→LHb glutamatergic axons were recorded in response to five footshocks one day before and one day after inescapable stress or no stress. A 2 (epoch) X 2 (day) x 2 (group) mixed ANOVA yielded a significant effect of epoch where shock-related activity was higher than baseline, F(1,8)=12.08, p < 0.0084. There were no main effects of day, group, or interactions. Instead, footshock significantly increased VTA→LHb axonal activity from baseline in the inescapable stress group both before inescapable stress, t(8)=-2.54,p=0.035, and after inescapable stress, t(8)=-3.665,p=0.006 (**Figure 1B-C**). In the no stress group, initial footshock did not significantly increase VTA→LHb axonal activity from baseline due to variability, but footshock significantly increased VTA→LHb axonal activity compared with baseline at the second exposure, t(8)= 4.216,p= 0.0059 (**Figure 1D-E**).

**Figure 1.**
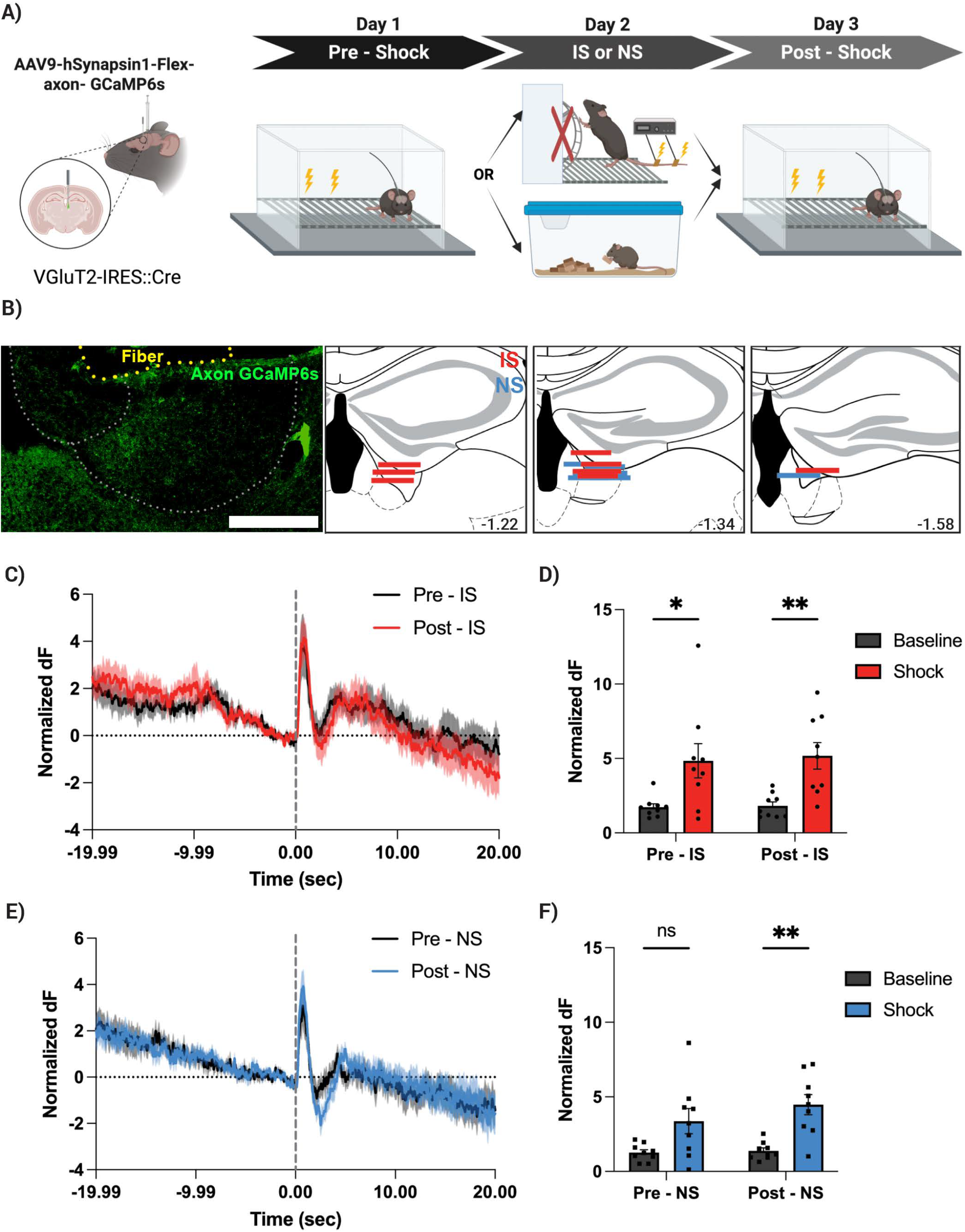
VTA inputs to LHb are activated by aversive events. **A**. Experimental setup. **B**. Example of axon GCaMP expression in LHb. Scale Bar is 200 µm. Colored lines represent localization of fiber optic. Numbers refer to mm from bregma. **C**. Average PETH of axon terminal calcium dynamics prior to and after inescapable stress. **D**. Quantifications of maximums for baseline and shock prior to and after inescapable stress. **E**. Average PETH of axon terminal calcium dynamics prior to and after no stress. **F**. Quantifications of maximums for baseline and shock prior to and after no stress. IS - inescapable stress, NS – no stress.

### 3.2 VTA glutamatergic activation results in LHb c-Fos expression following inescapable stress

We next assessed whether LHb neurons were differentially activated by VTA glutamatergic inputs in mice with or without prior inescapable stress experience (**Figure 2A**). VGluT2::Cre mice were injected in VTA with AAVs encoding Cre-dependent ChRmine-oScarlet and optic fibers were implanted dorsal to LHb. ChRmine stimulation of VTA inputs to LHb resulted in LHb c-Fos expression that did not differ between inescapable stress and no stress groups, Mann Whitney test z = -0.42, p = 0.7209. Collectively, VTA inputs to LHb were activated by aversive stimuli and their ability to drive LHb c-Fos is not modified by prior exposure to inescapable stress.

**Figure 2.**
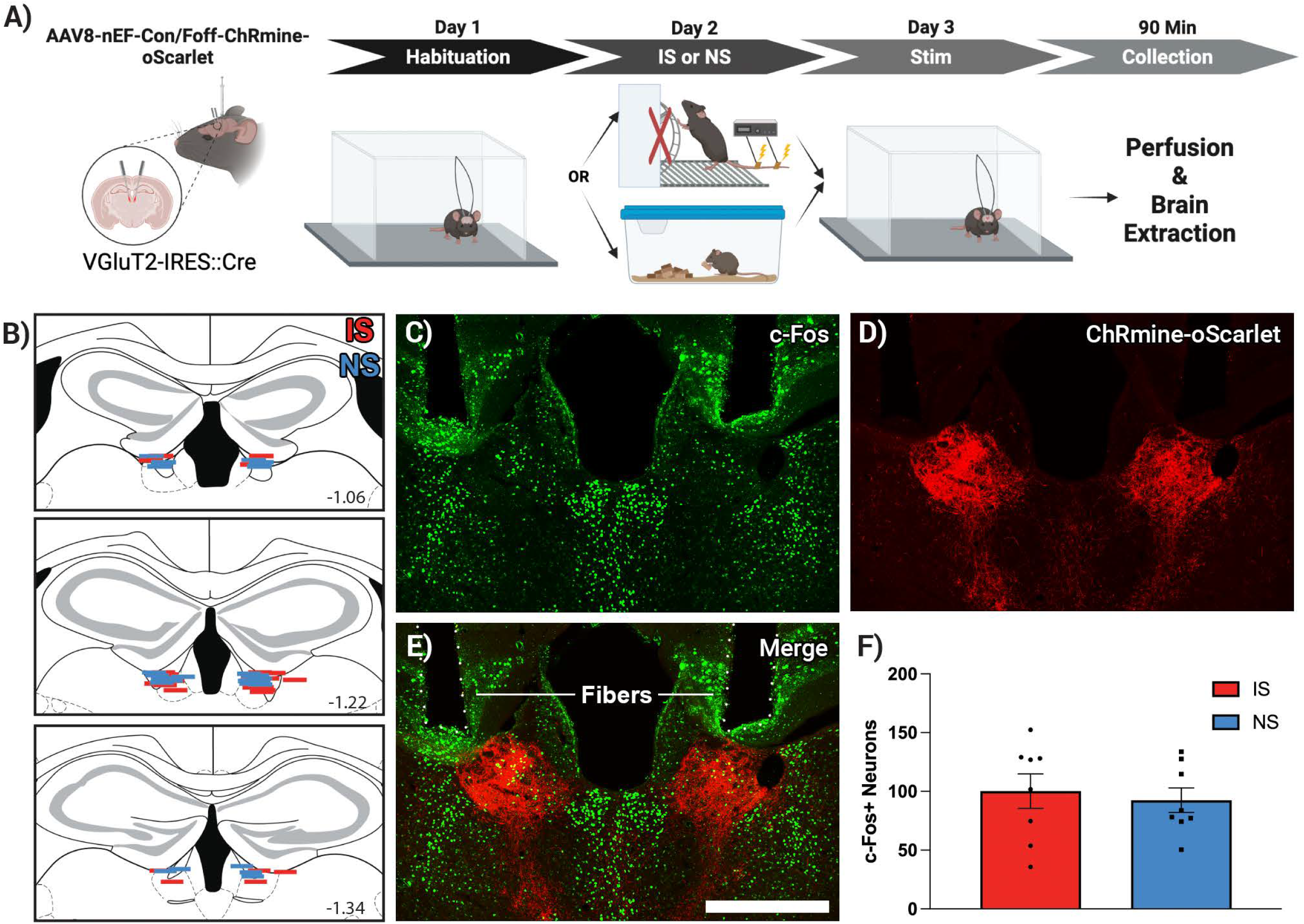
LHb c-Fos expression following IS or NS. **A**. experimental set up. **B**. Colored lines represent localization of fiber optic. Numbers refer to mm from bregma. **C**. Immunohistochemical labeling of c-Fos. Green neurons are c-Fos positive. Scale Bar is 500 µm. **D**. ChRmine-oScarlet expression in LHb. **E**. Merge c-Fos labeling and ChRmine-oScarlet expression. **F**. Quantification of c-Fos positive neurons following inescapable stress or no stress. IS - inescapable stress, NS – no stress.

### 3.3 VTA inputs to LHb are required for the consequences of inescapable stress

To test whether VTA→LHb glutamatergic axons regulate the behavioral consequences of inescapable stress, male and female VGluT2::Cre mice were injected in VTA with AAVs encoding Cre-dependent halorhodopsin or GFP and optic fibers were implanted dorsal to LHb (**Figure 3A**). To provide additional specificity on the projection, a group of VGluT2::Cre/VGaT::Flp mice were injected in VTA with AAVs encoding Cre and Flp dependent halorhodopsin and optic fibers were implanted dorsal to LHb to specifically target the glutamate and GABA co-transmitting pathway. Wildtype C57Bl6/J mice were used as escapable stress subjects and their tailshock stress durations and intensities were yoked to each inescapable stress group (GFP, Glu Halo, Glu/GABA Halo). For all inescapable stress mice, 589 nm light was delivered to LHb between tailshock onset and offset. Escapable stress mice progressively learned the wheel-turn response to terminate tailshock, F(8, 72) = 4.51,p = 0.0002, of which the fourth through the last block of trials showed significantly greater responding than the first block of ten trials (all p < 0.05) (**Figure 3C**). One day later, we assessed post-stress behaviors in a sex-specific fashion. In males, social exploration and fear-related behaviors was evaluated as previously described (McGovern et al., 2024). For social exploration, a 2 (social object) X 4 (group) mixed ANOVA yielded a significant interaction of social object and group, F(1, 8) = 42.07, p=0.0002. Sidak-adjusted multiple comparisons showed that escapable stress mice had a social preference by spending significantly more time with the social object than nonsocial object (p=0.0003) while inescapable stress GFP controls spent statistically indistinguishable times between objects (p= 0.0791). Mice that received inescapable stress and halorhodopsin inhibition of VTA glutamatergic inputs to LHb (p= 0.0140) or VTA glutamate/GABA inputs to LHb (p= 0.0016) each resulted in a statistically significant social preference (**Figure 3D**). Social preferences were confirmed in the social score, where positive numbers reflect more time with the social object and negative numbers reflect more time with the nonsocial object, relative to total exploration time. A between-subjects ANOVA yielded a significant effect of group, F(3, 27) = 5.643, p=0.0039. Sidak-adjusted multiple comparisons showed that inescapable stress GFP mice had lower social scores compared to escapable stress mice (p=0.0147), inescapable stress glutamate Halo mice (p= 0.0014), and inescapable stress glutamate/GABA Halo mice (p= 0.0424) (**Figure 3E**).

**Figure 3.**
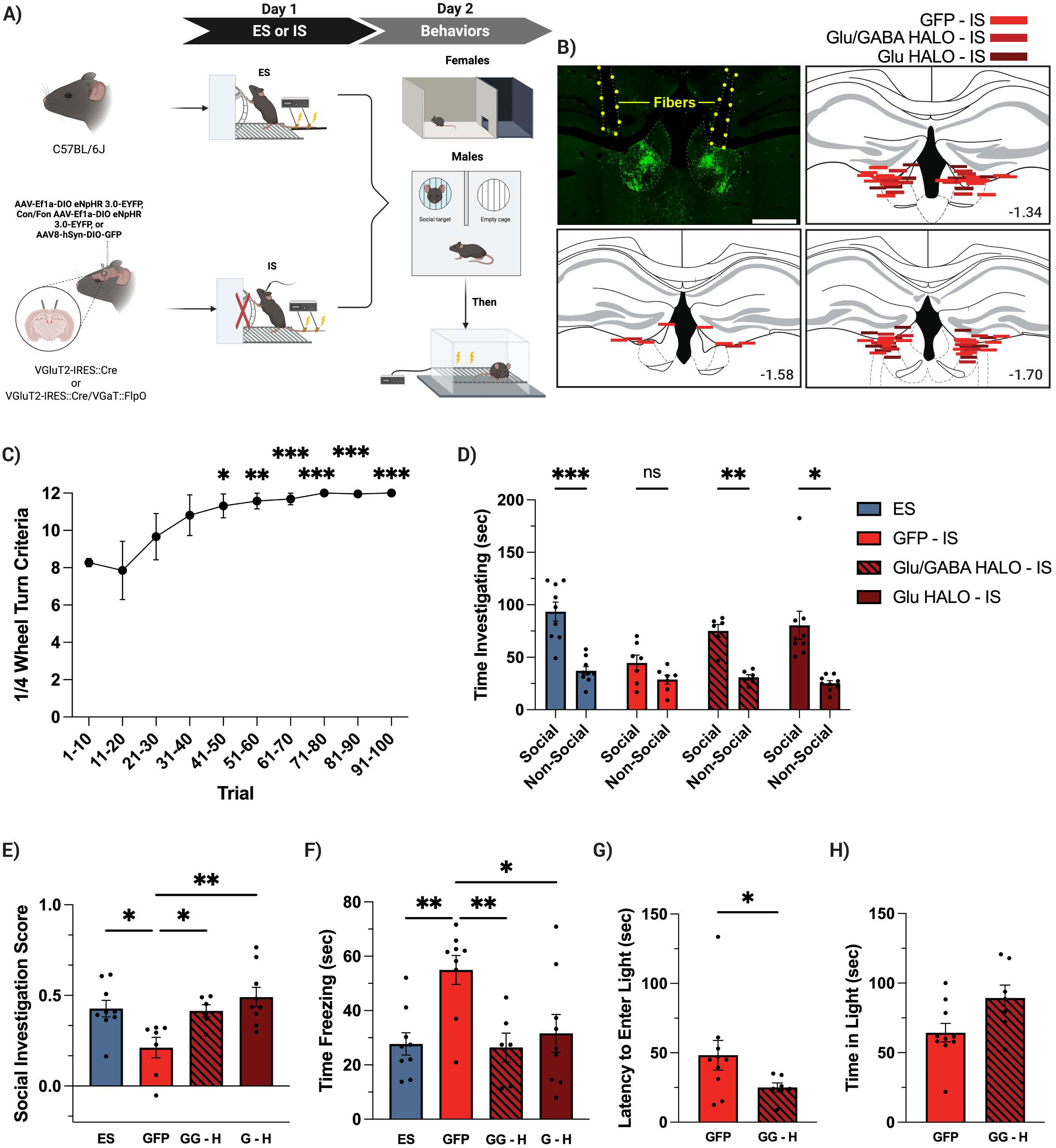
VTA glutamatergic axons in LHb are necessary for the social and nonsocial consequences of inescapable stress. **A**. Experimental setup. **B**. Example GFP expression in LHb. Scale Bar is 500 µm. Colored lines represent localization of fiber optic. Numbers refer to mm from bregma. **C**. Quarter wheel turns satisfying criterion to terminate tailshock over ten blocks of ten trials. Significance is compared to first block of trials. **D**. Time with social and nonsocial stimuli in male mice across groups. **E**. Social scores across groups in male mice. **F**. Time spent freezing in male mice across groups. **G**. Latency to enter the light side of the light dark box in female mice. **H**. Time spent in light side of the light dark box in female mice. ES – escapable stress; GFP – green fluorescent protein inescapable stress control group; G – H VTA glutamate → LHb halorhodopsin; GG – H VTA glutamate-GABA → LHb halorhodopsin. p < 0.05, ** p < 0.01, ns not significant.

We next evaluated fear-related behavior following to two footshocks in a novel context in males. A between-subjects ANOVA yielded a significant effect of group, F(3, 29) = 5.823, p=0.0031. Sidak-adjusted multiple comparisons showed that inescapable stress GFP mice froze significantly more than escapable stress mice (p= 0.0038), inescapable stress glutamate Halo mice (p= 0.0142), and inescapable stress glutamate/GABA Halo mice (p= 0.0068) (**Figure 3F**).

To determine if VTA inputs to LHb regulate the consequences of inescapable stress in female mice, female VGluT2::Cre/VGaT::Flp or VGluT2::Cre mice were injected in VTA with AAVs encoding Cre and Flp dependent halorhodopsin or GFP, respectively, and optic fibers were implanted dorsal to LHb. Both GFP and Halo inescapable stress groups received 589 nm light delivered to LHb between tailshock onset and offset. We previously showed that female mice show exploration-related deficits following inescapable stress and do not show social and fear-related behavior changes as male mice, but show changes in exploration within the light-dark box (McGovern et al., 2024). Therefore, one day after inescapable stress, females were examined in the light-dark box for exploration-related behaviors. We found that inescapable stress halorhodopsin mice had a significantly reduced latency to enter the light side of the light-dark box compared to inescapable stress GFP control mice, Mann-Whitney Test z=-2.051, p=0.0405 **(Figure 3G)**. However, there was no difference in time spent in the light compartment of the light-dark box between groups, z=-1.854, p=0.064 (**Figure 3H**).

## 4.0 DISCUSSION

The LHb is required for the consequences of inescapable stress (Amat et al., 2001; Dolzani et al., 2016). One source of input to LHb is from VTA VGluT2-expressing neurons, many of which co-transmit glutamate and GABA to LHb (Root et al., 2014b). Optogenetic activation of VTA glutamatergic axons in the LHb supports conditioned place aversion that is dependent on LHb glutamate receptors, suggesting an involvement of the VTA glutamatergic pathway to LHb in aversive processing (Root et al., 2014a; Lammel et al., 2015).

Chemogenetic inhibition of VTA glutamate neurons during inescapable stress blocked the consequences of inescapable stress in male and female mice (McGovern et al., 2024). However, the circuits by which VTA glutamate neurons regulate the consequences of inescapable stress are unknown. Here, halorhodopsin inhibition of VTA glutamatergic axons projecting to LHb, or halorhodopsin inhibition of VTA glutamate-GABA axons projecting to LHb during inescapable stress, blocked the consequences of inescapable stress in male and female mice. Thus, one source by which the LHb controls the consequences of inescapable stress is through glutamatergic inputs from VTA.

We previously found that about one-third of LHb neurons receiving glutamatergic synapses from the VTA were c-Fos activated following inescapable stress (McGovern et al., 2024). However, it was unknown whether VTA glutamate → LHb presynaptic axons or LHb neurons receiving VTA glutamatergic inputs were altered by prior inescapable stress. We found that VTA glutamate → LHb presynaptic axons were activated by footshock but their signaling of this aversive event was not altered by prior inescapable stress compared to mice with no prior stress experience. Further, ChRmine-stimulation of VTA glutamate → LHb axons resulted in statistically indistinguishable numbers of c-Fos expressing LHb neurons in mice with or without prior inescapable stress experience. It is possible that ChRmine stimulation resulted in a ceiling effect of c-Fos expressing neurons. However, because VTA glutamate-GABA neurons can scale their signaling of both rewarding and aversive stimuli (McGovern et al., 2024; McGovern et al., 2025), we interpret these results such that VTA glutamatergic inputs to LHb are involved in reporting salience of the immediate footshock stimulus which remained unchanged between pre and post stress conditions. Given that VTA glutamate neuron somatic calcium-related activity does not discriminate inescapable from escapable stress (McGovern et al., 2024), it is likely that VTA glutamatergic input to LHb coincides with other glutamatergic inputs related to stressor controllability that together regulate the consequences of inescapable stress.

While we recapitulated the chemogenetic rescue of inescapable stress-induced social and fear-related changes by more specific VTA→LHb glutamatergic axonal inhibition, females showed a partial rescue. Chemogenetic inhibition of all VTA glutamate neurons in females reduced the latency to enter the light side of the light dark box as well as increased the total time spent in the light side of the light dark box in inescapable stress mice (McGovern et al., 2024). Here, optogenetic inhibition of VTA glutamate-GABA axons in LHb reduced the latency to enter the light side of the light dark box but did not affect the total time spent in the light side of the light-dark box in female mice. It is possible that we were underpowered to detect this effect. Nevertheless, we previously observed that latency to enter the light side of the light dark box was sensitive to the controllability of the stressor while total time spent in the light side of the light dark box was not (McGovern et al., 2024). Given that a common feature of VTA glutamate neuron subtypes is their activation following aversive stimuli (Root et al., 2018; Barbano et al., 2020; Root et al., 2020; McGovern et al., 2021; Abdul et al., 2022; Barbano et al., 2024; Prevost et al., 2024; Warlow et al., 2024; McGovern et al., 2025; Prevost et al., 2025), it is also possible that the partial rescue of female post-stress behavior involves additional LHb pathways from the VTA (i.e., VGluT2+VGaT- or VGluT2+TH+ neurons (Root et al., 2014b)). A second possibility is the involvement of other VTA glutamatergic pathways. For example, halorhodopsin inhibition of all VTA glutamatergic axons in nucleus accumbens shell during repeated footshock blocked the development of behavioral despair (Qi et al., 2016). In addition, VTA glutamate neurons also project to bed nucleus of the stria medullaris (Root et al., 2020), which is required for the consequences of uncontrollable stress (Hammack et al., 2004).

The VTA→LHb glutamate circuits will also be modulated by their inputs. Of input circuits to VTA→LHb glutamatergic neurons that may be relevant to inescapable stress, we hypothesize the involvement of glutamatergic lateral hypothalamus neurons. Lateral hypothalamus VGluT2 neurons are critical regulators of VTA VGluT2 neuron function during defensive behavior (Barbano et al., 2020; Barbano et al., 2024), are required for the activation of VTA glutamate-GABA neurons in response to footshock (Prévost et al., 2024), and a driver of LHb’s influence on aversive processing (Stamatakis et al., 2016; Lecca et al., 2017; Lazaridis et al., 2019; Trusel et al., 2019; Rossi et al., 2021; Gu et al., 2023). Glutamatergic lateral hypothalamus neurons synapse onto VTA projecting LHb neurons (Poller et al., 2013), of which VTA projecting LHb neurons are potentiated by learned helplessness (Li et al., 2011), suggesting the possibility of reciprocal interactions between LH, VTA, and LHb during aversive processing that result in the consequences of inescapable stress.

Prior work has demonstrated that inescapable, but not escapable, stressors activate serotonergic neurons of the dorsal raphe (Grahn et al., 1999). This serotonergic activation is both necessary and sufficient for producing the behavioral sequelae of inescapable stress (Baratta et al., 2023). Upstream, dorsal raphe projecting LHb neurons are required for both the activation of dorsal raphe serotonin neurons by inescapable stress as well as the behavioral consequences of inescapable stress (Amat et al., 2001; Dolzani et al., 2016). Our results indicate that a glutamatergic mesohabenular pathway is required for the behavioral consequences of inescapable stress, but this pathway is not altered in presynaptic activity or postsynaptic influence on LHb following inescapable stress. We interpret these data such that the glutamatergic mesohabenular pathway may work together with other afferents toward a LHb tipping point that results in dorsal raphe serotonergic activation and the social and nonsocial sequelae of inescapable stress.

## Acknowledgements

Research supported in this publication was supported by the National Institutes of Health under award numbers F31 MH125569 (DJM), R01 MH050479 (MVB), R01 MH130576 (DHR/MVB), and R01 MH137472 (DHR). The content is solely the responsibility of the authors and does not necessarily represent the official views of the National Institutes of Health. The funders had no role in study design, data collection and analysis, decision to publish, or preparation of the manuscript. GraphPad Prism, Adobe Photoshop, and BioRender.com were used to generate figures and schematics.

## AUTHOR CONTRIBUTIONS

Conceptualization DJM, DHR, MVB

Data curation DJM, MD

Formal analysis DJM, MD

Funding acquisition DJM, DHR

Investigation DJM, MD, HM, HP

Supervision MVB, DHR

Visualization DJM, MD, DHR

Writing – original draft DJM

Writing – review and editing, MD, MVB, DHR

